# Temperature and nutrient conditions modify the effects of phenological shifts in predator-prey communities

**DOI:** 10.1101/2021.09.27.461998

**Authors:** V.H.W. Rudolf

## Abstract

While there is mounting evidence indicating that the relative timing of predator and prey phenologies shapes the outcome of trophic interactions, we still lack a comprehensive understanding of how important the environmental context (e.g. abiotic conditions) is for shaping this relationship. Environmental conditions not only frequently drive shifts in phenologies, but they can also affect the very same processes that mediate the effects of phenological shifts on species interactions. Thus, identifying how environmental conditions shape the effects of phenological shifts is key to predict community dynamics across a heterogenous landscape and how they will change with ongoing climate change in the future. Here I tested how environmental conditions shape effects of phenological shifts by experimentally manipulating temperature, nutrient availability, and relative phenologies in two predator-prey freshwater systems (mole salamander-bronze frog vs dragonfly larvae-leopard frog). This allowed me to (1) isolate the effect of phenological shifts and different environmental conditions, (2) determine how they interact, and (3) how consistent these patterns are across different species and environments. I found that delaying prey arrival dramatically increased predation rates, but these effects were contingent on environmental conditions and predator system. While both nutrient addition and warming significantly enhanced the effect of arrival time, their effect was qualitatively different: Nutrient addition enhanced the positive effect of early arrival while warming enhanced the negative effect of arriving late. Predator responses varied qualitatively across predator-prey systems. Only in the system with strong gape-limitation were predators (salamanders) significantly affected by prey arrival time and this effect varied with environmental context. Correlations between predator and prey demographic rates suggest that this was driven by shifts in initial predator-prey size ratios and a positive feedback between size-specific predation rates and predator growth rates. These results highlight the importance of accounting for temporal and spatial correlation of local environmental conditions and gape-limitation in predator-prey systems when predicting the effects of phenological shifts and climate change on predator-prey systems.

## Introduction

Phenology, the seasonal timing of life-history events, is a key force structuring species interactions. The relative timing of phenologies within a community determines what species and stages co-occur and thus can interact directly, when interactions start, and how long they last (Yang and Rudolf 2010). However, the relative timing of phenologies naturally vary across time and space (Carter et al. 2018, Rudolf 2018, Roslin et al. 2021) and is further altered by ongoing climate change (Parmesan and Yohe 2003, Visser and Both 2005, Cohen et al. 2018, Kharouba et al. 2018). While recent studies indicate that these changes in the relative timing of phenologies can alter the outcome of interactions and change long-term conditions for persistence and coexistence (Rudolf 2019), we are still lacking a general understanding of how important the environmental context (e.g. abiotic conditions) is for mediating the effects of phenological shifts. Yet environmental conditions vary across space and time (including climate change), and these differences are often (Visser and Holleman 2001, Durant et al. 2007, Dijkstra et al. 2011, Ovaskainen et al. 2013, Cohen et al. 2018), but not always (Roslin et al. 2021), the driver of phenological shifts. Thus, elucidating how the effects of phenological shifts vary across environmental conditions is not only essential to understand community dynamics across heterogeneous landscapes, but also key to predict how they will change in the future with ongoing climate change.

The potential for environmental conditions to modify the effects of phenological shifts becomes quickly clear when we focus on the link between phenologies and interactions. Phenological shifts can directly alter species interactions in at least two key ways: (i) by changing the temporal overlap of interacting species which determines their “interaction potential” (Carter et al. 2018) (i.e. how many individuals interact and for how long), and (ii) through shifts in per capita interaction strength (Rudolf 2019). Importantly, both mechanisms depend on growth and developmental rates of individuals. The duration of interactions between life-history stages (i.e. temporal overlap) generally decreases with higher growth and/or developmental rates because individuals transition to the next life history stage (phenophase) faster. Changes in per-capita effects driven by phenological shifts are frequently caused by concurrent shifts in size-ratios of interacting species: differences in arrival time allow early arrivers to grow and increase in relative size which determines per capita interaction strength (size-mediated priority effects) (Rasmussen et al. 2014). This suggests that any change in environmental conditions that influence the growth (and/or developmental) rates of species, such as temperature or nutrient availability, could also modify the consequences of phenological shifts for species interactions. Moreover, if these conditions have the same effect on growth rates we might also expect that they have the same qualitative effects on phenological shifts. If true, this would allow for general “rules of thumb” to predict what conditions strengthen or weaken effects of phenological shifts.

Despite the clear potential for environmental conditions to alter the effects of phenological shifts, this is rarely tested explicitly, and much remains unknown. Previous studies either examined phenological shifts only in one environmental context (e.g. Alford 1989, Nosaka et al. 2014, Rasmussen et al. 2014, Rasmussen and Rudolf 2016, Anderson et al. 2017), or used observational data (Durant et al. 2007, Visser and Gienapp 2019) for which the covariance of phenological shifts and environmental conditions make it inherently difficult to isolate individual and interactive effects of phenology vs. environment (Rafferty et al. 2013). The few recent experiments that manipulate phenologies either across different temperature or nutrient conditions seem to provide first support for context-dependent yet predictable effects of phenological shifts (Rudolf and Singh 2013, Rudolf 2018, Rudolf and McCrory 2018). However, these experiments only studied one environmental factor at a time and thus do not allow for direct comparisons of different environmental factors. Furthermore, they only focused on systems where species from the same trophic level compete for shared resources, and it is not straightforward to extrapolate their results to predator-prey systems. As a consequence, it remains unclear how different environmental conditions affect phenological shifts in predator-prey systems.

To understand the differences between competitive and predator-prey systems, let’s focus on two well-studied environmental factors: temperature and nutrient availability. With resource competition, interacting species experience the same temperature and the same nutrient levels and both are increasing growth and developmental rates. Thus, it is perhaps not surprising that both environmental factors appear to have qualitatively similar effects on phenological shifts in competitive systems (Rudolf and Singh 2013, Rudolf 2018, Rudolf and McCrory 2018). In contrast, predator and prey both experience the same temperatures, but they consume different resources and thus could respond differentially. For instance, an increase in primary productivity will directly benefit an herbivore, but not its specialized predator. The predator could still benefit indirectly if the available prey biomass eventually increases, but the response would be delayed, and increased growth rates of the prey would still likely reduce the time the prey is vulnerable to predation. A higher prey growth rate could be particularly important in systems with strongly gape-limited predators (e.g. predators that swallow prey whole) because it would allow prey to reach a size refuge and thus “escape” predation at an earlier stage (Wilbur 1988, Urban 2007). This suggests that nutrient availability and temperature could have qualitatively different effects on phenological shifts in predator-prey systems, and the effects could further depend on how gape-limited predators are.

Finally, we should not forget that temperature and nutrient availability also have different direct effects on other aspects of predator-prey interactions. Temperature directly affects size-specific predation rates, e.g. due to changes in attack rates and handling time (Uiterwaal and DeLong 2020). For instance, a moderate increase in temperature typically increases size-specific per-capita consumption rates of predators (Jara et al. 2019) and strengthens top-down control (Barton and Schmitz 2009, Shurin et al. 2012). In contrast, nutrient availability does not have this direct effect on per-capita predation rates. Indeed, increasing nutrient availability may instead indirectly decrease predation rates, e.g. by increasing availability of alternative prey (Chesson 1989, Rudolf 2008). Overall, this suggests that while temperature and nutrient availability both clearly have the potential to modify the consequences of phenological shifts in predator-prey systems, their individual effects could be qualitatively different compared to competitive systems and even vary across different predator-prey systems.

Here I take an experimental approach to test how environmental conditions influence the effects of phenological shifts on predator-prey interactions in two freshwater systems. Specifically, I experimentally manipulated the relative arrival time of a predator and its prey under different nutrient and temperature conditions. This allowed me to determine (1) how temperature and nutrient availability alter effects of phenological shifts, (2) whether this interactive effect qualitatively differs between both environmental factors, and (3) if their effects are independent or synergistic. Furthermore, I repeated the same experiment in two different predator-prey systems to determine (4) whether patterns are general or contingent on specific traits (e.g. gape-limitation) of predators and prey. Overall, results indicate that the phenological shifts affect demographic traits of both predator and prey, but the effects are modified by warming and nutrient availability and thus depend on the environmental context.

## Methods

### Study Species

I focused on two different predator-prey systems that are commonly found in fishless temporary ponds throughout the southwest of North America: (I) dragonfly larvae of the green darner *Anax junius* (predator) and tadpoles of the southern leopard frog *Rana* (*Litobathes*) *sphenocephala* (prey), and (II) larvae of the mole salamander *Ambystoma talpoideum* and tadpoles of the bronze frog *Rana* (*Lithobathes*) *clamitans*. Phenologies of these species vary naturally across years with changes in weather conditions. Because species respond differently to weather conditions, changes in these conditions result in concurrent changes in the onset of species interactions (Root et al. 2003, Saenz et al. 2006, Heino et al. 2009, Todd et al. 2011, Carter et al. 2018). Furthermore, ponds naturally differ in temperature regimes and nutrient input (e.g. due to variation in canopy cover) (Skelly et al. 2002), creating considerable spatial heterogeneity in these conditions.

The two predator-prey systems differ in many aspects from each other. Larvae of the dragonfly *A. junius* and the salamander *A. talpoideum* are both major predators of tadpoles in fishless pond communities (Wellborn et al. 1996, Wilbur 1997), but they differ in their morphology, ecology, and behavior. Specialized mouthparts (two opposing thorn-like structures) allow dragonflies to capture and consume prey much larger than themselves. In contrast, the suction feeding of salamanders limits them to consume prey that is smaller than their mouth’s diameter (Urban 2008). This strong gape-limitation allows tadpoles to “escape” predation from salamander by reaching a size-refuge at a certain predator/prey size ratio (Caldwell et al. 1980, Urban 2008). This is not the case with dragonfly predators in our system, although successful attack rates still typically decline with smaller predator/prey size ratios (Caldwell et al. 1980). Indeed, while monitoring the ponds we observed dragonflies that attacked and ultimately killed tadpoles many times larger than the dragonflies themselves.

The two tadpole species also differ in their growth and developmental rate and phenology at our field sites in South East Texas. Southern leopard frogs are active foragers with high growth rates and a mean larval period (hatching to metamorphosis) of 90 days. In contrast, bronze frog tadpoles are much less active and mostly hide in the substrate and leave litter. As a consequence, they have lower growth rates and a much longer larval period, lasting up to 22 months, and frequently overwinter in ponds before reaching metamorphosis in spring.

### Experimental design

Both experiments shared the exact same factorial design which crossed 3 tadpole phenology (“arrival”) treatments (tadpole addition 0 days, +10 days, or +20 days after predator addition) with 2 nutrient (ambient vs. enriched) and 2 temperature (ambient vs. heated) treatments, resulting in a total of 3 × 2 × 2 = 12 treatments (**Fig. 1**). In the first experiment (with dragonfly predators) each treatment was replicated 5 times. In the second experiment (with salamander predators) treatments were replicated 4 times because of logistic constraints. A mistake during early tadpole addition during setup of salamander experiment resulted in uneven replication in low (ambient) nutrient treatments with 3 replicates for early (day 0) and 5 replicates for intermediate (+10 days) treatments (see supplement for details).

**Figure 1:**
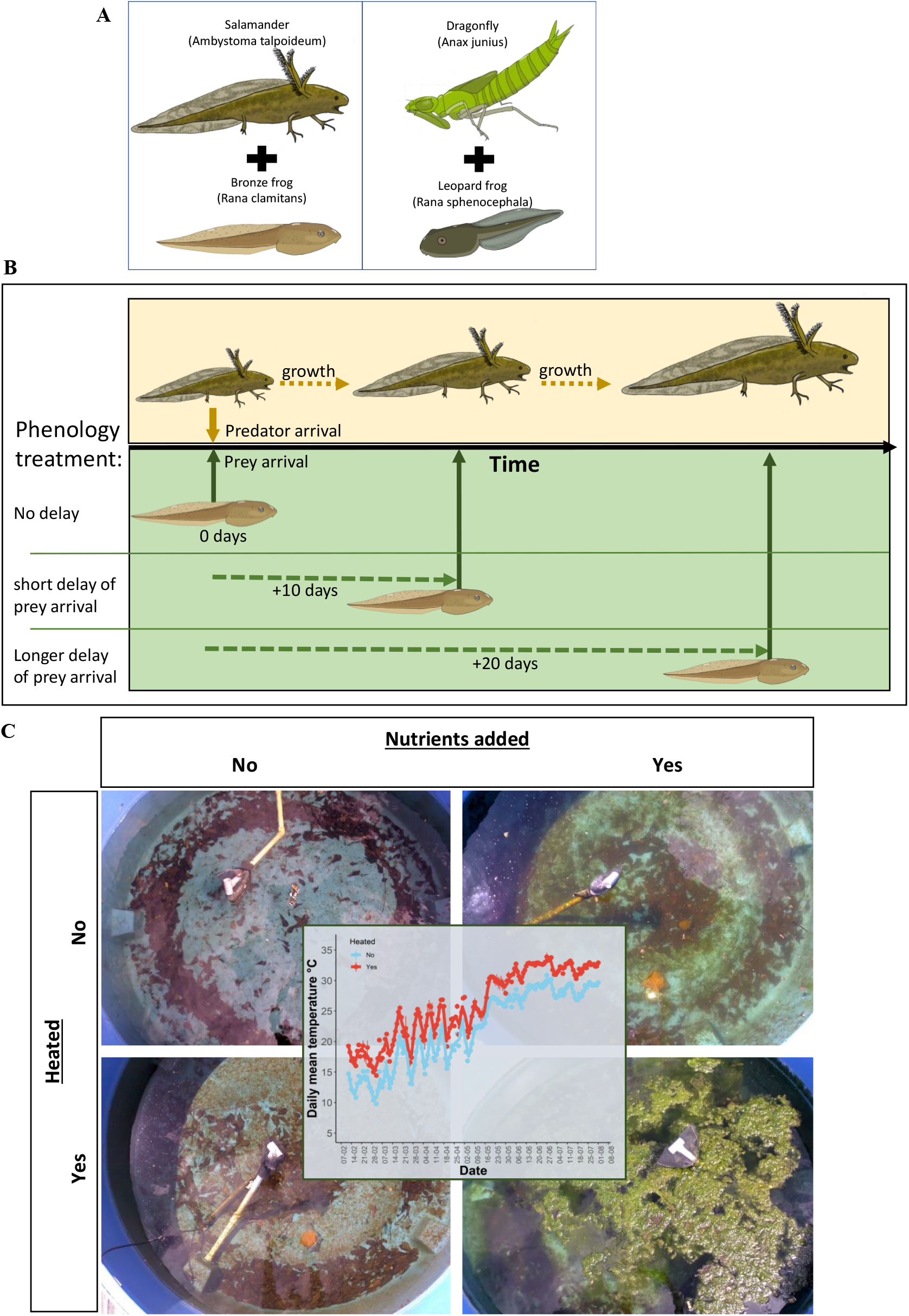
Experimental setup and study system. **(A)** The two predator-prey systems used in this study. Salamander larvae preying on bronze frogs, and dragonfly larvae preying on leopard frogs. Both systems differ in various ways from each other (see methods for details) including how much gape limitation plays a role in predation. (**B**) Phenology treatments manipulated relative arrival time of prey, which were either added on the same day as predators or arrival was delayed by 10 or 20 days relative to predator to test for effects of phenological shifts. These phenology treatments were repeated across (**C**) different environmental conditions, by manipulating temperature (heated vs. ambient) or availability of limiting nutrients in experimental mesocosms. Pictures show representative examples of how treatments influenced mesocosm communities. Insert shows temporal fluctuations in daily temperatures for heated vs. ambient mesocosms.

To delay prey arrival (hatching) for the different phenology treatments, I transferred all collected egg clutches to a climate-controlled environmental chamber maintained at 4°C to slow down development. One week before a given arrival date I transferred a subset of clutches to another environmental chamber set at 24°C to accelerate development and hatch tadpoles. Previous experiments indicate that this method can be successfully used to delay hatching by up to 25 days without measurable effects on tadpole performance (Rudolf and Singh 2013, Rudolf 2018). It also assured that tadpoles from all additions were within the same developmental stage and had the same size at introduction. For each introduction, I used 3-4 randomly selected egg clutches and distributed tadpoles from all clutches evenly across replicates. Each replicate in both experiments received 200 tadpoles of the respective prey species. These starting densities are well within the natural range of both species we observe in our study region.

In the dragonfly experiment, each mesocosm received three small *A. junius* larvae of equal size (mean head width (HW) = 3.296mm, range = 2.983-3.392mm). Mesocosms in the salamander experiment each received five small *A. talpoideum* larva. Due to natural variation in size, salamanders were visually divided into three size classes and each size class was evenly distributed among replicates to ensure similar mean and variation in predator size across all replicates (mean HW: 3.655 mm (2.607-4.616), snout-vent length SVL: 11.340 mm (7.769-15.035). The respective size range of both predators reflects the natural size range of predator populations when tadpoles hatch in natural ponds. The densities of both predators are at the lower end of natural densities in our study area for the size classes. Differences in initial densities between both predators reflect species-specific differences in natural densities and predation rates.

I terminated the dragonfly predator experiment after 189 days (January 29^th^ - August 5^th^) and the salamander experiment after 100 days (February 9^th^ - May 15^th^). This difference in duration reflects natural differences in developmental times of predator species: mean emergence time was 91.9 days (range 62-123 days) for dragonflies and 59.9 for salamanders (range: 49-88 days). The additional time also allowed me to better capture differences in the timing of metamorphosis of *R. sphenocephala* (> 80% of individuals reached metamorphosis in all arrival treatments at end of the experiment), while *R. clamitans* was not close to reaching metamorphosis due to its naturally slower developmental rate.

### Mesocosm setup & maintenance

I conducted experiments in mesocosms consisting of 1,000 L plastic cattle tanks set up outside at the South Campus Research Facility of Rice University. Mesocosms were evenly spaced by 0.5m and filled with dechlorinated well water two weeks before tadpole addition. I covered each mesocosm with 60% shade cloth to reduce unwanted colonization by other amphibians or predators. To establish natural conditions, I added 1L of dried leaf litter and 500mL of concentrated zooplankton and pond water, collected from fishless ponds where all species occur naturally. This setup followed well-established protocols and allowed me to create replicate communities that mimicked key aspects of temporary ponds used by all species (Wilbur 1997, Rudolf and Rasmussen 2013b, Rudolf and Rasmussen 2013a).

I manipulated temperature by placing one 300-Watt submersible heater at the center of each mesocosm one week before the start of the experiment and wrapped each mesocosm with insulation. This setup allowed heated replicates to follow the same natural daily and seasonal temperature fluctuations as ambient mesocosms but elevated mean temperatures by ∼4.1°C and 4.7°C in salamander and dragonfly predator experiment respectively (supplement **Fig. 1, S1**). I assigned heating treatments spatially so that no two heated mesocosms were next to each other. Temperatures were monitored every half hour with iButton® temperature loggers that were submerged in a subset of heated and ambient mesocosms.

Like most freshwater systems, temporary pond communities are frequently limited by nutrients, especially nitrogen and phosphorus (Schindler 1977, Mischler et al. 2014). Thus, for the nutrient treatment, I either left mesocosms at ambient levels or added nitrogen and phosphorus (Nitrogen:7.31g/100L, Phosphorous 0.30618g/1000L). Nutrient additions were based on similar experiments (Kratina et al. 2012) and pilot studies and ensured a significant increase in algae growth without the risk of creating large bacterial blooms that can cause anoxic conditions. In the first experiment (with dragonfly predators) I added nutrients twice, one week before tadpole addition and then again after tadpole addition. Since I observed a decline in phosphorus throughout the first experiment, I repeated nutrient additions in the second experiment 38 days and 94 days after tadpole addition to maintaining elevated nutrient levels throughout the experiment. Both nutrient additions were successful in significantly elevating nutrients and primary production (see supplement **Fig. 2, S5**).

### Response variables

#### Primary producers

I measured periphyton (benthic algae) density (primary food source of tadpoles) throughout the experiment. I quantified periphyton density by floating 3 microscope glass slides per mesocosm for 7 days and extracting chlorophyll *a* from periphyton scraped off from both sides of each slide following standard protocols (Eaton et al. 2005). Slides were replaced every 1-2 weeks throughout the experiment.

#### Predator and prey

I measured predator size (SVL and HW) at each tadpole introduction to quantifying differences in initial predator size across tadpole introductions using photographs and ImageJ. Predators, especially dragonflies were very difficult to subsample without draining mesocosms. Thus, to minimize and standardize disturbance caused by these subsamples, I spend a fixed time (15 minutes) per mesocosm which assured that I caught at least one predator per mesocosm. In addition, I measured 20 tadpoles per mesocosm 13-18 days after the final tadpole introduction to quantifying initial tadpole growth rates.

I monitored mesocosms daily and collected all predators and prey that reached metamorphosis. For tadpoles, day of metamorphosis was defined as the emergence of at least one forelimb, and all metamorphs were transferred to the lab and weighed after full tail absorption. Salamander metamorphosis was defined by absorption of external gills. Emerging dragonflies were easy to count but very difficult to catch alive, preventing us from collecting sufficient body size data for a full analysis. I calculated growth rates and developmental for all species except *R. clamitans* which did not reach metamorphosis, and final body size/mass for all but *A. junius*. At the end of the experiment, I destructively sampled all mesocosms and collected any remaining tadpoles and predators. Surviving tadpoles were photographed to measure the snout-vent length (SVL). All procedures followed recommended guidelines of the Animal Welfare Act and were approved by Institutional Animal Care and Use Committee (IACUC protocol A13101101)

### Statistical analyses

While both experiments had the exact same setup and design, using different species and carrying them out in different years inherently results in some natural variation in abiotic and biotic conditions and what response variables could be quantified. Thus, they should be considered as two separate experiments that address the same questions and I analyzed them separately. Any interpretation of differences across experiments should keep these caveats in mind. However, because both experiments used the same experimental design and asked the same questions, comparing qualitative relationships across different predator-prey systems still provides valuable insights into how sensitive results are to differences in species’ life histories and helps identify general patterns.

#### Predators

Initial predator-prey size ratios can drive effects of phenological shifts. I used generalized linear mixed models (GLMM) to test how predator size changed with each prey introduction and if this relationship was affected by nutrient addition and temperature, using predator size at each introduction as dependent variable and arrival time (= days since the start of the experiment), heating, and nutrient addition treatments as predictors, and mesocosm identity as a random factor to account for non-independence of repeated observations. Note that for the dragonfly experiment we only measured predator size in mesocosms without prey present, while I measured predators in all mesocosms in the salamander experiment. However, prey arrival order did not affect the initial size of salamanders. Therefore, I pooled those treatments for final analysis. Finally, I used general linear models (GLM) with final predator size (for salamander), time to emergence, or survival as the response variable, and heating, nutrient, and arrival treatment and all possible interactions as fixed effects.

#### Prey

I used GLMs to analyze treatment effects on per-capita size, daily (dry) biomass production (final prey dry mass/days since prey addition), and mortality rates (number of prey that died /days since prey addition). Using daily mortality rate and biomass production allowed me to correct for inherent differences in the time individuals spent in the experiment across arrival treatments and thus allows for a direct comparison across arrival treatments. Tadpoles and metamorphs were all converted to dry mass using established mass-length relationships and dry mass was summed across metamorphs and tadpoles within a replicate.

All analyses were carried out in R using the “lme4” package for GLM analyses and the “car” package to obtain significance values. I used a binomial error distribution for salamander survival and Gaussian distributed error for all other analyses. Because of unbalanced replication in the salamander experiment, P-values are based on type III statistics while all other P-values are based on type II unless noted otherwise. The corresponding code and data is freely available online at dryad: (*will be added once accepted for publication*)

## Results

### Changes in initial predator-prey size ratios across prey arrival times

How predator size (measured by head width HW) differed across prey arrival times was contingent on environmental conditions in both predator systems (significant heating x time interaction, **Table 1, Fig. 2)**. Predator size remained largely unchanged between the last two prey arrival times (day 10 vs. 20) under ambient conditions, but it increased significantly in heated systems (**Fig. 2**). Average predator size also increased in heated relative to ambient treatments (**Fig 2, Table 1**). In contrast, nutrient addition did not affect the size of dragonfly predators, but it enhanced the positive effect of warming on salamander growth rate and size (**Fig. 2, Table 1**). As a consequence, when prey arrival was delayed by 20 days, salamander predators were up to 68% larger in heated communities with added nutrients (head width 6.24mm) compared to predators in systems with ambient temperature and nutrient conditions (head width 4.26 mm).

**Table 1:**
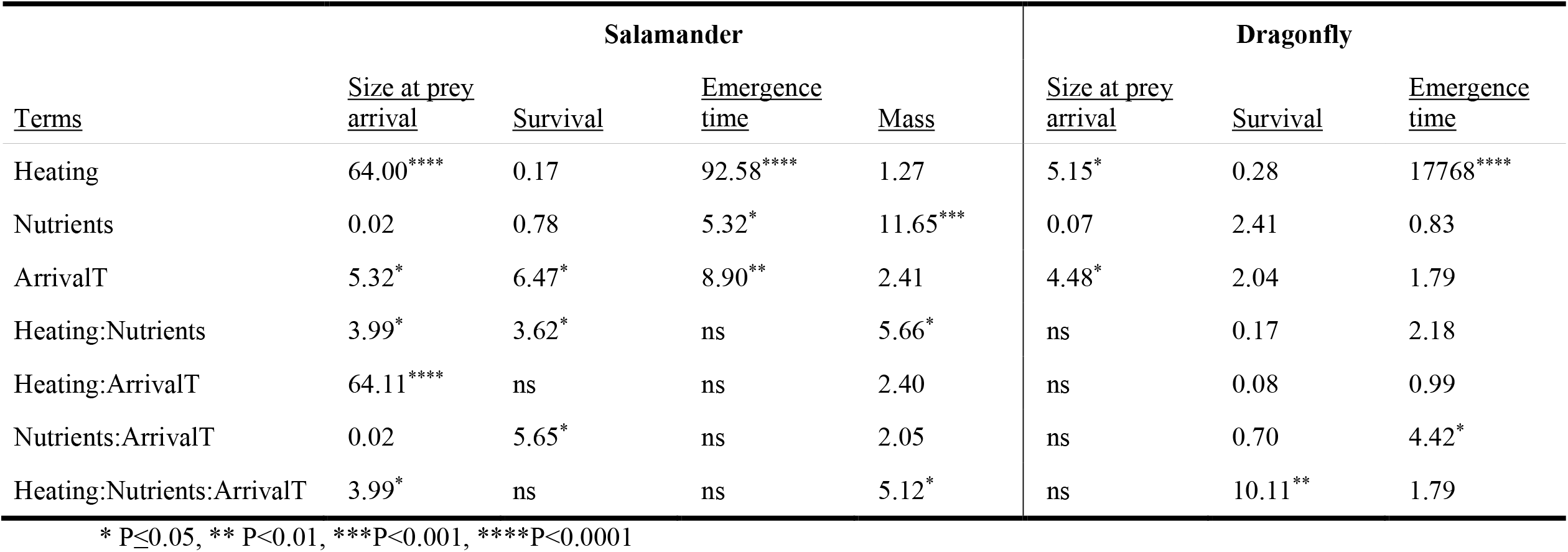
Effects of delay in prey arrival time (ArrivalT), heating, and nutrient addition treatments on predator demographic traits. Values indicate Wald Chisquare statistics for a given demographic trait. All values show type III statistics for salamander and size at prey arrival for dragonflies and type II for remaining dragonfly survival and emergence time. ns indicates that interactions were not significant (P>0.05) and dropped for final model for type III statistics to facilitate interpretation of main effects.

**Figure 2:**
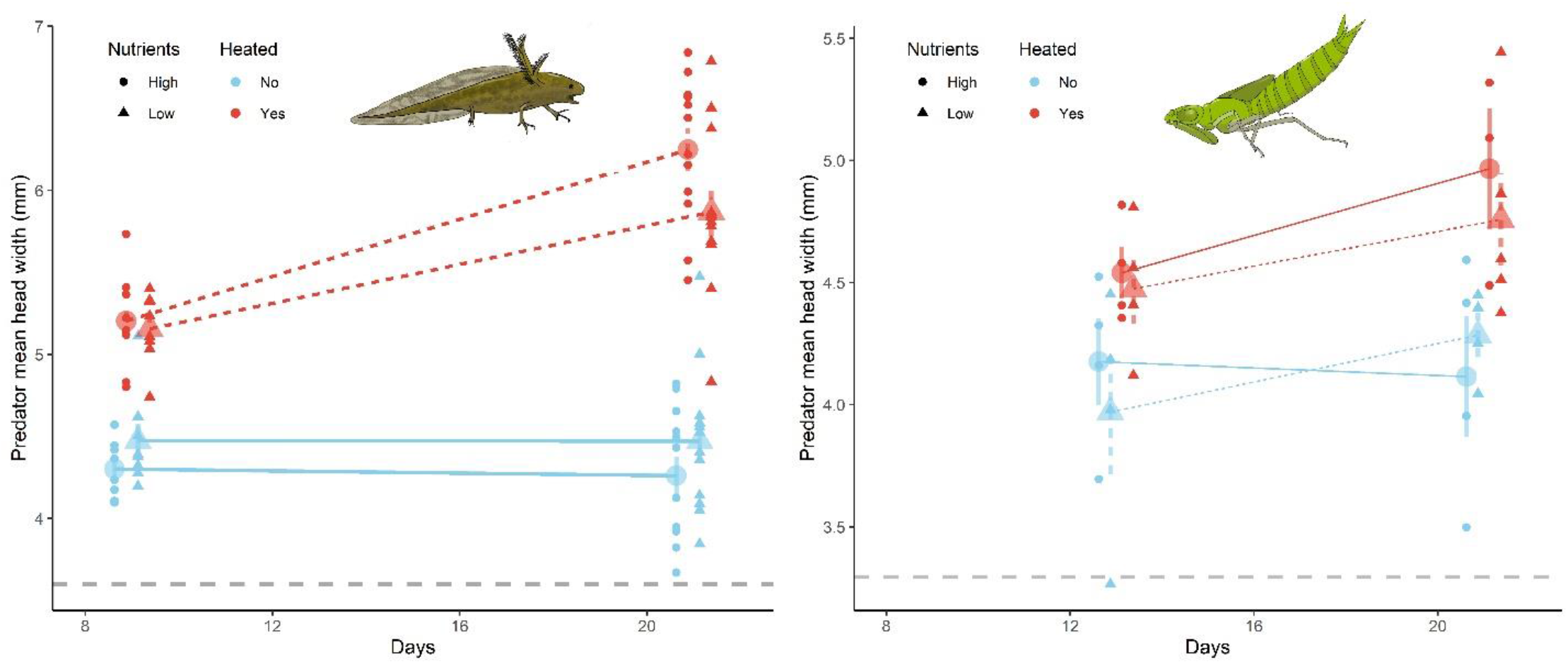
Change in predator size with delay (10 and 20 days) in prey arrival across different temperature (heated) and nutrient conditions. Days indicate the number of days that have passed after predators were added to the experiment. Large symbols indicate treatment mean ±1 SE, small symbols indicate individual mesocosm means. Symbols of different treatments are offset horizontally for a given sample day for visual clarity. Dashed grey horizontal lines indicate the respective mean size at the start of the experiment (day 0). Note that for logistic reasons, the first sample was taken a few days later for dragonflies, and samples size was smaller for dragonfly experiment because subsamples were restricted to replicates without tadpoles, while this was not the case for salamander experiment (tadpole presence did not significantly affect predator size, see methods for more details).

### Predator survival, development & growth

Predator survival was high for both predator species, especially for salamander (mean survival = 89.4%). Predator survival significantly decreased when prey were introduced later (**Table 1**) for salamander but not dragonfly systems. There was some indication of nutrient treatment affecting arrival in salamander but this was solely driven by a single low survival outlier and not significant after removing the outlier. Dragonfly survival was driven by three-way interaction (**Table 1**). Survival increased with delay in arrival time in systems at ambient temperature and high nutrients or heated and low nutrients, while it remained largely constant or even declined in the other two treatment combinations. Note that random invasion of dragonfly predators in some tanks (indicated by the number of survivors > number of added focal individuals) prevented any exact estimates of dragonfly survival.

Both predator species had significantly higher developmental rates resulting in ∼29-31% shorter emergence time in heated treatments (dragonflies: 103 days vs 78.4, salamander 67.6 days vs 52.4 days, **Fig. 3**). Salamander, but not dragonfly development was also significantly faster under high nutrient treatments and when prey arrived later, although this effect was much smaller than the warming effect (**Table 1, Fig. 3**). Salamander mass at metamorphosis was determined by interactions of all three treatments, prey arrival time, heating, and nutrients (three-way interaction, P = 0.0237) (**Table 1, Fig. 3**): their mass increased the later prey arrived, but this increase was largest (68%) in heated treatments with high nutrients where predators (mass increased from 889.5mg to 1,495.4mg). Together, these results indicate that a delay in prey arrival time increased developmental and growth rates of salamander predators, and this relationship was strengthened by nutrient addition and warming in a given community.

**Figure 3:**
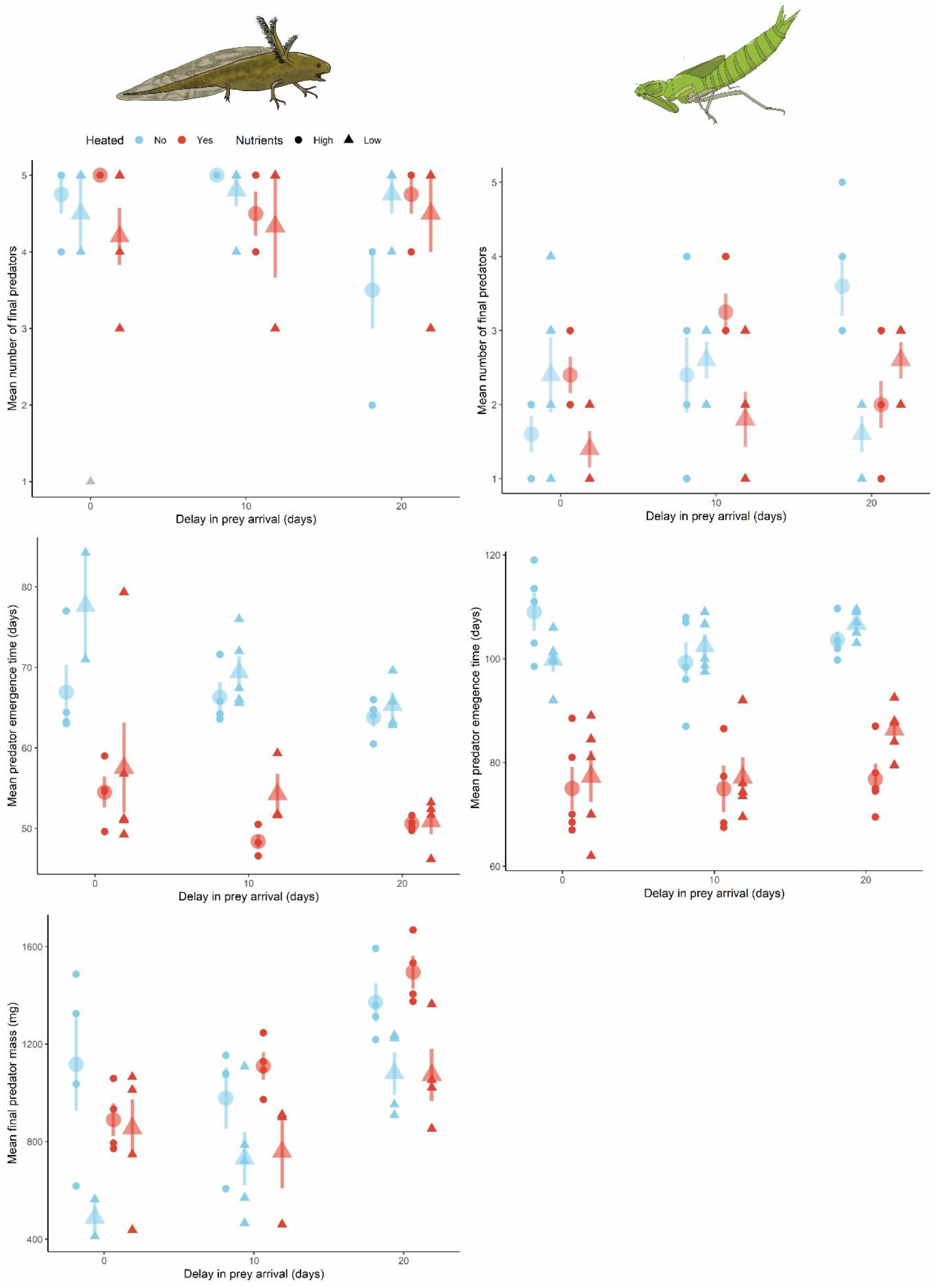
Effects of delay in prey arrival time on predator demographic traits across different nutrient and temperature conditions. Large symbols indicate mean ±1 SE, small symbols indicate mesocosm means. Symbols of different treatments are offset horizontally for a given arrival day treatment for visual clarity. The grey point in the top right panel indicates an outlier and was not included in the mean or final statistical analysis (see results).

### Prey response

As expected, prey that arrived later were significantly smaller during the early part of the experiment (33-38 days after start) under ambient conditions (**supplemental Table S2)**. In contrast, size differences between early vs. later arriving prey were much smaller (or even absent) in heated communities (**Fig. S2**). Together with effects on predator size, this confirms that arrival treatments modified predator-prey size ratios with delay in prey arrival time and this was further modified by differences in temperature regimes. However, growth rates showed the opposite pattern and increased significantly with delay in prey arrival, indicating that experimentally delaying arrival did not negatively affect early growth rates (**supplemental Table S2)**. In leopard frogs (with dragonfly predators) nutrient addition also significantly increased size and growth rates (**supplemental Table S2)**. This suggests that prey size differences created by differences in relative arrival should decline over time, especially in heated communities and high nutrient levels.

Delaying prey arrival significantly increased prey mortality rates in both predator-prey systems (**Table 4, Fig 4**): a 20-day delay in prey arrival increased mortality rates on average by 1.2 to 2.2 times in dragonfly and salamander predator systems respectively. Warming and nutrient addition both significantly affected mortality rates (**Table 2**), but their effects differed qualitatively from each other and between both predator-prey systems. In both predator-prey systems, nutrient addition significantly reduced prey mortality, but this effect was strongest with early prey arrival and declined significantly the later prey arrived (**Table 2, Fig. 4**). As a consequence, a delay in arrival time had a stronger effect (steeper increase in mortality, **Fig. 4**) in treatments with added nutrients, but this interaction was only significant in experiments with dragonfly predators (**Table 2**). Heating significantly increased prey mortality in both experiments (**Table 2**). However, in salamander experiments, this temperature effect was much stronger when prey arrival was delayed, essentially enhancing the negative effect of delay in prey arrival time (steeper increase in mortality) (ambient: early: 136 vs late: 100 survivors; heated: early: 103 vs late: 13.5 survivors) (**Fig. 4**). Thus, nutrient addition and warming both enhanced the effects of delaying prey arrival, but nutrients enhanced the positive effect of early prey arrival while warming enhanced the negative effect of late arrival.

**Figure 4:**
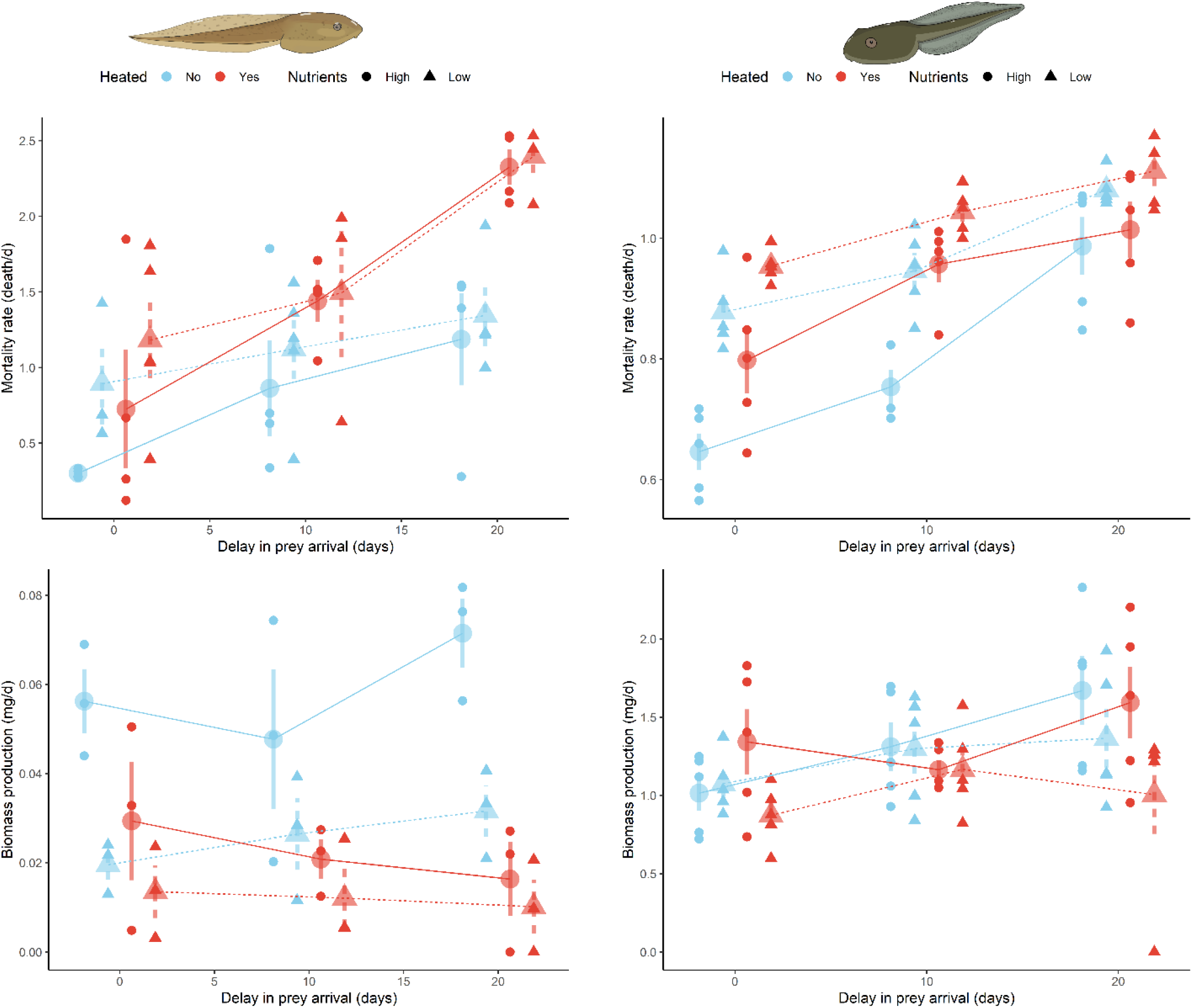
Effect of delay in prey arrival time on prey demographic traits across different nutrient and temperature conditions. Mortality rate and biomass production indicate a daily change in prey survival and total dry biomass within a given treatment. Large symbols indicate mean ±1 SE, small symbols indicate mesocosm means. Symbols of different treatments are offset horizontally for a given arrival day treatment for visual clarity.

**Table 2:** Effects of delay in prey arrival time (ArrivalT), heating, and nutrient addition treatments on prey demographic traits. Biomass production indicates total biomass produced per time averaged across the duration of the experiment. Values indicate Wald Chisquare statistics for a given demographic trait. All values show type III statistics for salamander and size at prey arrival for dragonflies and type II for remaining dragonfly survival and emergence time. ns indicates that interactions were not significant (p>0.05) and dropped for the final model for type III statistics to facilitate interpretation of main effects.

| <u>Terms</u> | **Bronze frog**<br>**(with salamander predator)** |  | **Leopard frog**<br>**(with dragonfly predator)** |  |
| --- | --- | --- | --- | --- |
|  | <u>Mortality rate</u> | <u>Biomass</u><br><u>production</u> | <u>Mortality rate</u> | <u>Biomass</u><br><u>production</u> |
| Heating | 2.10 | **8.73\*\*** | **26.07\*\*\*\*** | 1.03 |
| Nutrients | **4.17\*** | **25.25\*\*\*\*** | **55.61\*\*\*\*** | **5.09\*** |
| ArrivalT | **8.07\*\*** | 2.96 | **95.48\*\*\*\*** | **7.84\*\*** |
| Heating:Nutrients | ns | **5.92\*** | 2.37 | 1.89 |
| Heating:ArrivalT | **4.46\*** | **3.79\*** | 3.31 | 1.40 |
| Nutrients:ArrivalT | ns | ns | **4.39\*** | 1.04 |
| Heating:Nutrients:ArrivalT | ns | ns | 0.71 | 0.27 |

\* $p \leq 0.05$ , \*\* $p < 0.01$ , \*\*\* $p < 0.001$ , \*\*\*\* $p < 0.0001$

Total prey biomass production was on average significantly higher with added nutrients in both experiments and lower in heated treatments in the salamander experiment (**Table 2**). In contrast to mortality, biomass moderately increased with delay in prey arrival in the dragonfly predator system. This relationship was driven by a very strong compensatory growth response in surviving individuals; the per-capita mass of surviving prey increased in all treatments with delay in prey arrival time by up to three times in high nutrient treatments (**supplemental Fig. S3**). In salamander predator systems, the arrival effect was contingent on the heating treatment, with a positive relationship in ambient systems and opposite (decline with arrival time) in heated communities. The decline in heated tanks occurred because the strong compensatory growth of individuals could not overcome the even stronger increase in mortality rate in heated systems. Finally, in the dragonfly experiment where prey developed much faster, a large proportion of prey completed metamorphosis, with shorter development time in either higher temperatures or nutrients and delay in arrival (**supplemental Fig. S3**).

### Predator-prey feedbacks

Predator size (head width) at prey introduction (indicating differences in initial predator/prey size ratio) was strongly positively related to prey survival, explaining 27% and 34% of the total variation in prey survival in salamander and dragonfly predator systems respectively (with salamander: F1,48 = 55.5, P< 0.0001, with dragonfly: F_1,58_ = 13.19, P=0.0006, **Fig. 5**). Furthermore, final salamander mass was positively (F_1,45_ = 5.0 P=0.03) and developmental time of both predators was negatively correlated (salamander: F_1,45_ = 18.5 P<0.0001, dragonfly: F_1,56_ = 13.19, P= 0.0006) with prey mortality (supplement **Fig. S4**), suggesting a feedback between consumed prey and predator growth and development.

**Figure 5:**
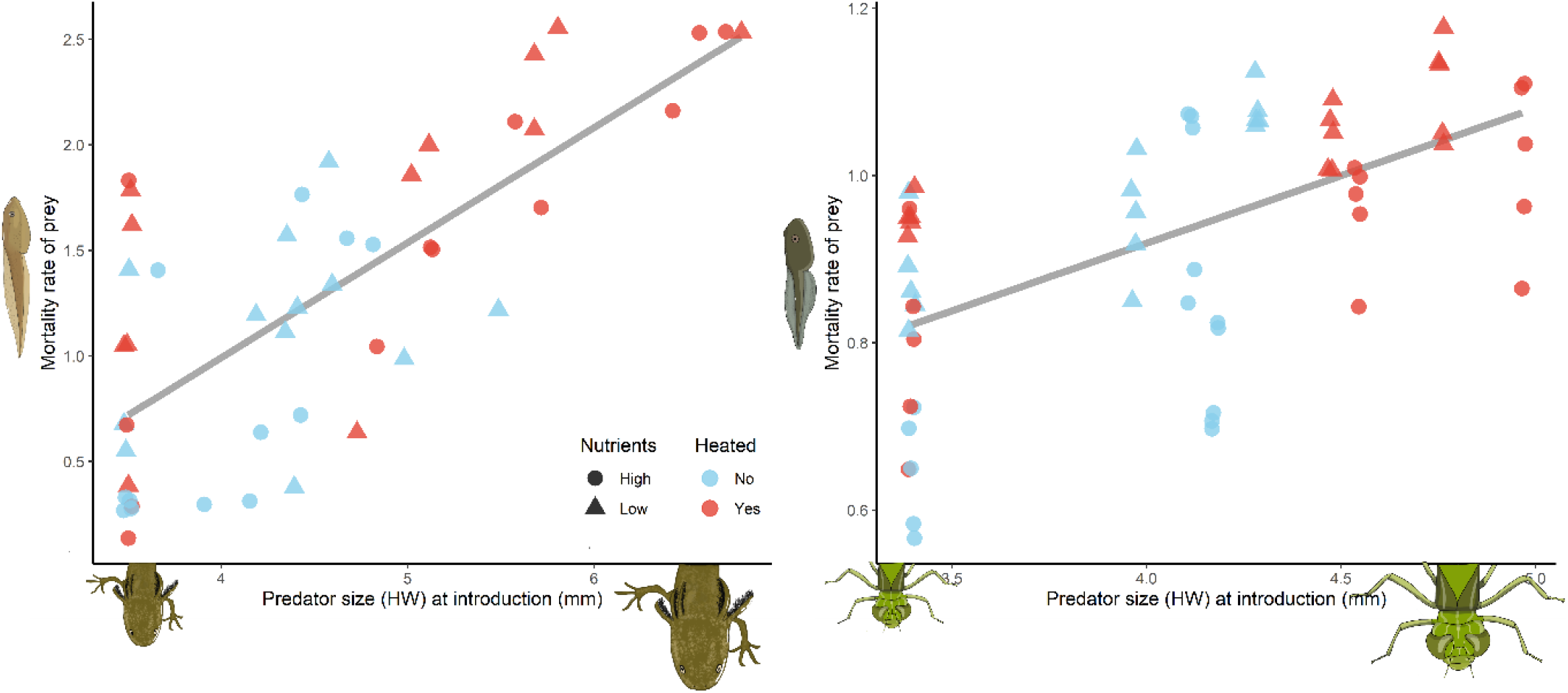
Relationship between predator size at prey introduction and prey survival. (A) salamander –bronze frog system, (B) dragonfly-leopard frog system. Grey lines indicate significant linear relationships. Symbols indicate individual replicates in a given treatment and indicate the mean size (measured as head width) of predators. Note differences in y-axis scaling between panels. Points are jittered by 0.01 for clarity to avoid overlapping data points.

## Discussion

### Phenological shifts alter predator-prey interactions

The relative timing of predator and prey phenologies can play a key role in shaping predator-prey systems. Previous research has largely focused on the concept of trophic (phenological) match/mismatch (Cushing 1969, Visser and Gienapp 2019, Kharouba and Wolkovich 2020). The trophic mismatch concept focuses on the temporal overlap of peak prey availability and peak energetic demands of predators (e.g. during reproduction)(Kharouba and Wolkovich 2020). While intuitively appealing, trophic match/mismatches are notoriously difficult to proof and explicit experimental tests are “extremely rare” (Visser and Gienapp 2019, Kharouba and Wolkovich 2020). Furthermore, this approach typically neglects that phenological shifts can also modify per-capita interaction strength (Rudolf 2019).

Using an experimental approach, I showed that shifts in phenologies can significantly affect both prey and predator populations. Several lines of evidence indicate that the observed effects of phenological shifts were driven by an increase in per-capita predation rates with delay in prey arrival. Consistent with previous studies (Rudolf and Singh 2013, Rasmussen and Rudolf 2016, Rudolf 2018, Rudolf and McCrory 2018, Carter and Rudolf 2019), I found no evidence that experimentally delaying prey hatching negatively affects prey performance. Indeed, late-arriving prey even grew and developed faster than early arriving prey, likely because their key food resources (periphyton) increased during that period. However, the increase in prey mortality with delay in its arrival time is consistent with an increase in per capita predation rates. Delaying prey arrival increased prey mortality, which was correlated with increased growth and developmental rates of predators. An increase in predation also explains why prey per-capita mass and developmental rates increased: predation reduced the density of prey, which reduced intraspecific competition in the prey and allowed prey to grow and develop faster. Similar plastic responses have been observed in other studies (Anderson et al. 2017, Carter and Rudolf 2019) and explain why prey biomass even increased with delay in prey arrival time.

The increase in predation rates with delay in prey arrival is consistent with size-mediated priority effects. Arriving earlier than their prey allows predators to grow to a larger size when interactions are initiated, which in turn should increase per-capita predation rates. Furthermore, since prey typically grow faster than predators, this size advantage of predators can also prolong the time prey are within a vulnerable size range of gape-limited predators. Consistent with this expectations, the size of predators at the time of prey introduction was a significant predictor of prey mortality and explained ∼30% of the variation in prey mortality. Size-mediated priority effects are known to play important role in mediating effects of phenological shifts in competitive systems (Rudolf and Singh 2013, Rudolf 2018, Rudolf and McCrory 2018, Blackford et al. 2020), with important consequences for long-term dynamics (Rudolf 2019). However, they are rarely considered in the predator-prey or trophic mismatch literature (Visser and Both 2005, Visser and Gienapp 2019, Kharouba and Wolkovich 2020). Yet, such changes in per-capita effects are likely to be common, especially when interactions occur among growing predators and prey (Wilbur 1988, Urban 2007, Yang and Rudolf 2010, Nosaka et al. 2014, Rasmussen and Rudolf 2016).

### The role of predator and prey traits in mediating phenological shifts

Predator-prey systems can differ in many ways from each other (e.g. growth rates, per-capita predation rates, predator and prey behavior, gape-limitation, etc.) and these differences could alter the effects of phenological shifts and relative importance of the environmental context. The two different systems I used here showed some remarkably similar patterns but also highlighted some key differences. Only salamander predators were strongly affected by prey arrival and its interaction with environmental conditions, while dragonfly predators only responded to changes in temperature regimes. This difference could at least partly be driven by the fact that salamanders are much more gape-limited. Size measurements indicate that most prey reached a size refuge from salamander predation (i.e. body width > salamander mouth diameter) at some point during the experiment. When prey arrive at the same time as predators, this likely happened early when predators were still small, decoupling prey performance and predator traits (e.g. no or weak correlation of predator mass and development rats and prey mortality, supplement). In contrast, when salamanders were relatively larger when prey arrived later, they were able to consume prey for longer (aided by a corresponding increase in predator growth rates). In contrast, dragonflies could always attack and kill prey, even when the prey was much larger. The dragonfly predator-prey systems thus should be much less sensitive to variation in initial predator-prey size ratio or relative growth rates. This helps explain why dragonfly predators were not affected by prey arrival time.

These results suggest that some effects of phenological shifts appear to be consistent across predator-prey systems, while others depend on the details of predator and prey traits. To date, we are missing studies that are specifically aimed to link traits of interacting species to the effects of phenological shifts. Yet identifying general mechanisms that link species traits to effects of phenological shifts is key to gain a comprehensive and predictive understanding of seasonal dynamics of communities and how they will be affected by future climate change. The results presented here provide an important step towards achieving this goal, but much more work is needed in a diversity of systems to address this gap in our knowledge.

### Environmental context mediates effects of phenological shifts

Environmental conditions frequently differ across systems and often co-vary with phenological shifts (Benard 2015, Cohen et al. 2018). I found that differences in environmental conditions can modify the effects of phenological shifts in predator-prey systems and in some rare instances multiple environmental factors can even interact with each other. However, warming and nutrient additions had qualitatively different effects. Warmer conditions increased and high nutrient levels dampened the negative effect of late arrival for prey (i.e. steeper increase in mortality with delay in arrival time). Furthermore, the effect of warming was stronger in the salamander predator system, while the effect of nutrients was stronger in the dragonfly system.

The interaction of warming and phenological shifts is again consistent with size-mediated priority effects. Warming had by far the strongest effect on initial predator size and increased predator size with delay in prey arrival relative to ambient conditions. Since predation rates are typically positively correlated with predator/prey size ratios (Urban 2007), this helps explain why delaying prey arrival time was associated with a steeper increase in prey mortality compared to ambient conditions. This size-mediated priority effect could also have been further enhanced by an increase in size-specific predation rates under warmer conditions (Jara et al. 2019).

In contrast to warming, nutrient addition enhanced the positive effects of early arrival. Nutrient addition had little or only minor effects on initial predator growth rates and predator sizes did not differ across prey arrival times. However, nutrient addition did increase initial and final growth rates as well the developmental rates of prey. This contrasting effect of nutrients on predator vs. prey reflects the simple fact that they consume different resources: prey (but not predators) consume periphyton, which was increased by nutrient addition (see supplement). It is also possible that differences in nutrient availability altered prey (or predator) behavior and thereby changed predation rates. Elucidating the relative contribution of different potential mechanisms was beyond the scope of this study, but the results presented here suggest that additional and previously overlooked factors (i.e. beyond size-mediated priority effects) plaid an important role. The results highlight the complex and dynamic interaction between environmental conditions and phenological shifts and the need for further research to understand how general these patterns are and what other mechanisms are involved.

In the broader context of climate change, the results also suggest that climate-mediated shifts in phenologies or temperature patterns (Benard 2015, Cohen et al. 2018) depend on how both are correlated and local conditions. For instance, a delay in prey arrival would have a much more negative effect on prey survival if it is correlated with an increase in temperature. Similarly, a delay in prey arrival is likely to heave weaker effects in systems with high nutrient (resource) availability for the prey. These results highlight that phenological shifts need to be considered in the respective environmental context of a system, and how they are correlated with shifts in environmental conditions.

## Data Availability

*All data will be made publicly available with corresponding code for statistical analysis on dryad with publication*

## Acknowledgments

I would like to thank Amber Roman, Zach Costa, Rebecca Maher, Beineng Zhan, Jane Giang, Lilly Stockset, Anecia Gentles, Bianca Romado, Elaine Shen and Avery Twitchik-Hayne for help with field and lab work, and Amy Dunham and Tom Miller for help in developing the ideas in the manuscript. This work was in part supported by funding from NSF award DEB-1256860 and DEB-1655626.

## References

Alford, R. A. 1989. Variation in predator phenology affects predator performance and prey community composition. Ecology 70:206–219.

Anderson, T. L., F. E. Rowland, and R. D. Semlitsch. 2017. Variation in phenology and density differentially affects predator–prey interactions between salamanders. Oecologia 185:475–486.

Barton, B. T., and O. J. Schmitz. 2009. Experimental warming transforms multiple predator effects in a grassland food web. Ecology Letters 12:1317–1325.

Benard, M. F. 2015. Warmer winters reduce frog fecundity and shift breeding phenology, which consequently alters larval development and metamorphic timing. Global Change Biology 21:1058–1065.

Blackford, C., R. M. Germain, and B. Gilbert. 2020. Species Differences in Phenology Shape Coexistence. The American Naturalist 195:E168–E180.

Caldwell, J., J. Thorp, and T. Jervey. 1980. Predator-prey relationships among larval dragonflies, salamanders, and frogs. Oecologia 46:285–289.

Carter, S. K., and V. H. W. Rudolf. 2019. Shifts in phenological mean and synchrony interact to shape competitive outcomes. Ecology 100:e02826.

Carter, S. K., D. Saenz, and V. H. Rudolf. 2018. Shifts in phenological distributions reshape interaction potential in natural communities. Ecology Letters 21:1143–1151.

Chesson, J. 1989. The effect of alternative prey on the functional response of Notonecta hoffmani. Ecology 70:1227–1235.

Cohen, J. M., M. J. Lajeunesse, and J. R. Rohr. 2018. A global synthesis of animal phenological responses to climate change. Nature Climate Change 8:224–228.

Cushing, D. 1969. The regularity of the spawning season of some fishes. ICES Journal of Marine 33:81–92.

Dijkstra, J. A., E. L. Westerman, and L. G. Harris. 2011. The effects of climate change on species composition, succession and phenology: a case study. Global Change Biology 17:2360–2369.

Durant, J. M., et al. 2007. Climate and the match or mismatch between predator requirements and resource availability. Climate research 33:271–283.

Eaton, A. D., et al. 2005. Standard methods for examination of water and wastewater. American Public Health Association, American Water Works Association, Water Environment Federation.

Heino, J., R. Virkkala, and H. Toivonen. 2009. Climate change and freshwater biodiversity: detected patterns, future trends and adaptations in northern regions. Biological Review 84:39–54.

Jara, F. G., et al. 2019. Warming-induced shifts in amphibian phenology and behavior lead to altered predator–prey dynamics. Oecologia 189:803–813.

Kharouba, H. M., et al. 2018. Global shifts in the phenological synchrony of species interactions over recent decades. Proceedings of the National Academy of 115:5211–5216.

Kharouba, H. M., and E. M. Wolkovich. 2020. Disconnects between ecological theory and data in phenological mismatch research. Nature Climate Change 10:406–415.

Kratina, P., et al. 2012. Warming modifies trophic cascades and eutrophication in experimental freshwater communities. Ecology 93:1421–1430.

Mischler, J. A., P. G. Taylor, and A. R. Townsend. 2014. Nitrogen Limitation of Pond Ecosystems on the Plains of Eastern Colorado. PLOS ONE 9:e95757.

Nosaka, M., N. Katayama, and O. Kishida. 2014. Feedback between size balance and consumption strongly affects the consequences of hatching phenology in size-dependent predator–prey interactions. Oikos 124:225–234.

Ovaskainen, O., et al. 2013. Community-level phenological response to climate change. Proceedings of the National Academy of 110:13434–13439.

Parmesan, C., and G. Yohe. 2003. A globally coherent fingerprint of climate change impacts across natural systems. Nature 421:37–42.

Rafferty, N. E., et al. 2013. Phenological overlap of interacting species in a changing climate: an assessment of available approaches. Ecology and Evolution 3:3183–3193.

Rasmussen, N. L., and V. H. W. Rudolf. 2016. Individual and combined effects of two types of phenological shifts on predator–prey interactions. Ecology 97:3414–3421.

Rasmussen, N. L., B. G. Van Allen, and V. H. W. Rudolf. 2014. Linking phenological shifts to species interactions through size-mediated priority effects. Journal of Animal Ecology 83:1206–1215.

Root, T. L., et al. 2003. Fingerprints of global warming on wild animals and plants. Nature 421:57–60.

Roslin, T., et al. 2021. Phenological shifts of abiotic events, producers and consumers across a continent. Nature Climate Change 11:241–248.

Rudolf, V. H. W. 2008. Impact of cannibalism on predator-prey dynamics: Size-structured interactions and apparent mutualism. Ecology 89:1650–1660.

Rudolf, V. H. W. 2018. Nonlinear effects of phenological shifts link interannual variation to species interactions. Journal of Animal Ecology 87:1395–1406.

Rudolf, V. H. W. 2019. The role of seasonal timing and phenological shifts for species coexistence. Ecology Letters 22:1324–1338.

Rudolf, V. H. W., and S. McCrory. 2018. Resource limitation alters effects of phenological shifts on inter-specific competition. Oecologia 188:515–523.

Rudolf, V. H. W., and N. L. Rasmussen. 2013a. Ontogenetic functional diversity: Size-structure of a keystone predator drives functioning of a complex ecosystem. Ecology 94:1046–1056.

Rudolf, V. H. W., and N. L. Rasmussen. 2013b. Population structure determines functional differences among species and ecosystem processes. Nature Communications 4:2318.

Rudolf, V. H. W., and M. Singh. 2013. Disentangling climate change effects on species interactions: effects of temperature, phenological shifts, and body size. Oecologia 173:1043–1052.

Saenz, D., et al. 2006. Abiotic correlates of anuran calling phenology: The importance of rain, temperature, and season. Herpetological Monographs:64–82.

Schindler, D. W. 1977. Evolution of phosphorus limitation in lakes. Science 195:260–262.

Shurin, J. B., et al. 2012. Warming shifts top-down and bottom-up control of pond food web structure and function. Philosophical Transactions of the Royal Society B-Biological Sciences 367:3008–3017.

Skelly, D. K., L. K. Freidenburg, and J. M. Kiesecker. 2002. Forest canopy and the performance of larval amphibians. Ecology 83:983–992.

Todd, B. D., et al. 2011. Climate change correlates with rapid delays and advancements in reproductive timing in an amphibian community. Proceedings of the Royal Society B: Biological Sciences 278:2191–2197.

Uiterwaal, S. F., and J. P. DeLong. 2020. Functional responses are maximized at intermediate temperatures. Ecology 101:e02975.

Urban, M. 2007. Predator size and phenology shape prey survival in temporary ponds. Oecologia 154:571–580.

Urban, M. C. 2008. Salamander evolution across a latitudinal cline in gape-limited predation risk. Oikos 117:1037–1049.

Visser, M. E., and C. Both. 2005. Shifts in phenology due to global climate change: the need for a yardstick. Proceedings of the Royal Society of London Series B-Biological Sciences 272:2561–2569.

Visser, M. E., and P. Gienapp. 2019. Evolutionary and demographic consequences of phenological mismatches. Nature ecology & evolution 3:879–885.

Visser, M. E., and L. J. M. Holleman. 2001. Warmer springs disrupt the synchrony of oak and winter moth phenology. Proceedings of the Royal Society of London Series B-Biological Sciences 268:289–294.

Wellborn, G. A., D. K. Skelly, and E. E. Werner. 1996. Mechanisms creating community structure across a freshwater habitat gradient. Annual Review of Ecology and 27:337–363.

Wilbur, H. M. 1988. Interactions between growing predators and growing prey. Pages 157-172 in B. Ebenman and L. Persson, editors. Size-structured populations - ecology and evolution. Springer Verlag, Berlin.

Wilbur, H. M. 1997. Experimental ecology of food webs: Complex systems in temporary ponds. Ecology 78:2279–2302.

Yang, L. H., and V. H. W. Rudolf. 2010. Phenology, ontogeny, and the effects of climate change on the timing of species interactions. Ecology Letters 13:1–10.

