## supplement for "Temperature and nutrient conditions modify the effects of phenological shifts in predator-prey communities"

**Supplement for: Temperature and nutrient conditions alter consequences of phenological shifts in predatory-prey communities**

Contains: Table S1, S2, Figures S1-S

**Table S1**: Treatments and corresponding number of replications (reps.) in two experimental systems: dragonfly – leopard frog vs. salamander – bronze frog. Note that because of one mistake in early introduction treatment (heated & nutrient added), replication had to be adjusted in two other treatments as mesocosms already had assigned temperature and nutrient treatments established.

| Heated | Nutrients | Prey arrival time | Dragonfly predator reps. | Salamander predator reps. |
|---|---|---|---|---|
| N | N | 0 | 5 | 4 |
| N | N | 10 | 5 | 4 |
| N | N | 20 | 5 | 4 |
| N | Y | 0 | 5 | 3 |
| N | Y | 10 | 5 | 5 |
| N | Y | 20 | 5 | 4 |
| Y | N | 0 | 5 | 4 |
| Y | N | 10 | 5 | 4 |
| Y | N | 20 | 5 | 4 |
| Y | Y | 0 | 5 | 5 |
| Y | Y | 10 | 5 | 3 |
| Y | Y | 20 | 5 | 4 |

**Figure S1: Temperature over time in mesocosms that were either not heated (ambient) or heated experimentally.** Individual points represent daily average temperatures across mesocosms. Note that heated and ambient mesocosms show exact same natural variation in temperature over time and only differ in mean temperature. (A) Salamander- bronze frog experiment, (B) dragonfly-leopard experiment. Note differences in x axis scales.

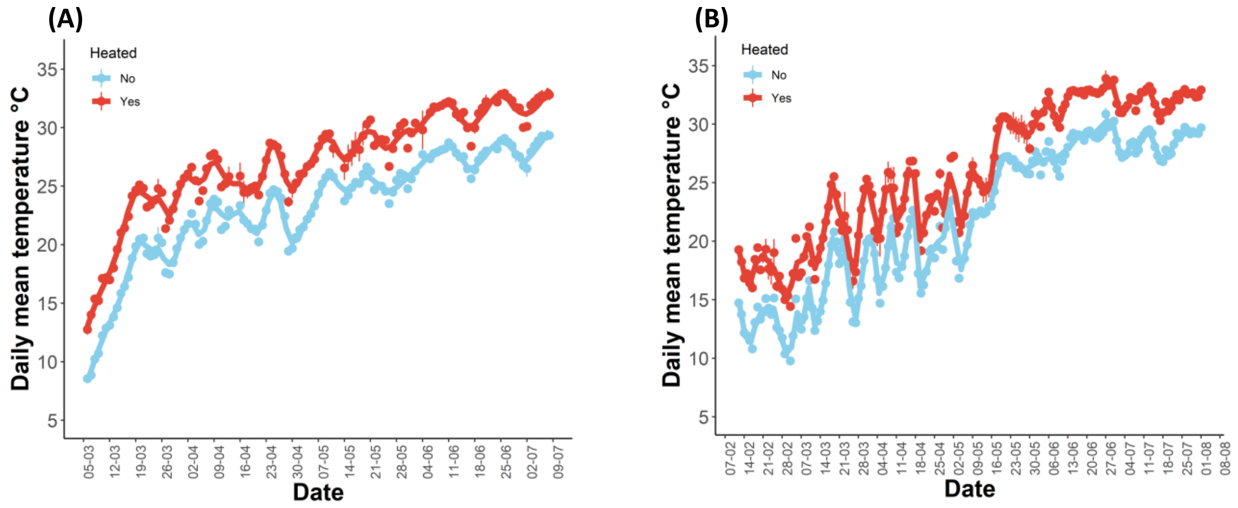

**Table S2:** Treatment effects on prey size (SVL) and growth rate 2-3 weeks after final introduction (specifically 33 and 38 days after first tadpole addition for dragonfly and salamander predator system respectively). Values indicate Wald Chisquare statistics for a given demographic trait. All values show type III statistics for bronze frog and type II for leopard frogs. ns indicates that interactions were not significant (p>0.05) and dropped for final model for type III statistics to facilitate interpretation of main effects.

| | **Bronze Frog (with salamander predator)** | | **Leopard Frog (with dragonfly predator)** | |
|---|---|---|---|---|
| Terms | **Size** | **Growth rate** | **Size** | **Growth rate** |
| Heated | 2.21 | 0.56 | 41.52**** | 80.30**** |
| Nutrients | 0.47 | 0.14 | 57.43**** | 68.05**** |
| ArrivalT | 57.30**** | 6.72** | 162.43**** | 116.04**** |
| Heated:Nutrients | 3.05 | 2.51 | 10.03** | 15.39*** |
| Heated:ArrivalT | 32.50**** | 89.46**** | 2.45 | 23.15**** |
| Nutrients:ArrivalT | 1.14 | 0.55 | 8.48** | 0.67 |
| Heated:Nutrients:ArrivalT | ns | ns | 0.04 | 1.00 |

* p≤0.05, ** p<0.01, ***p<0.001, ****p<0.0001

**Figure S2.** Size and growth rates of prey 33-38 days after start of experiment. (A, C) bronze frog (with salamander predator), (B, D) leopard frog (with dragonfly predator). Large symbols indicate mean ±1 SE, small symbols indicate mesocosm means. Symbols of different treatments are offset horizontally for a given arrival day treatment for visual clarity. Note differences in y axis scale between species.

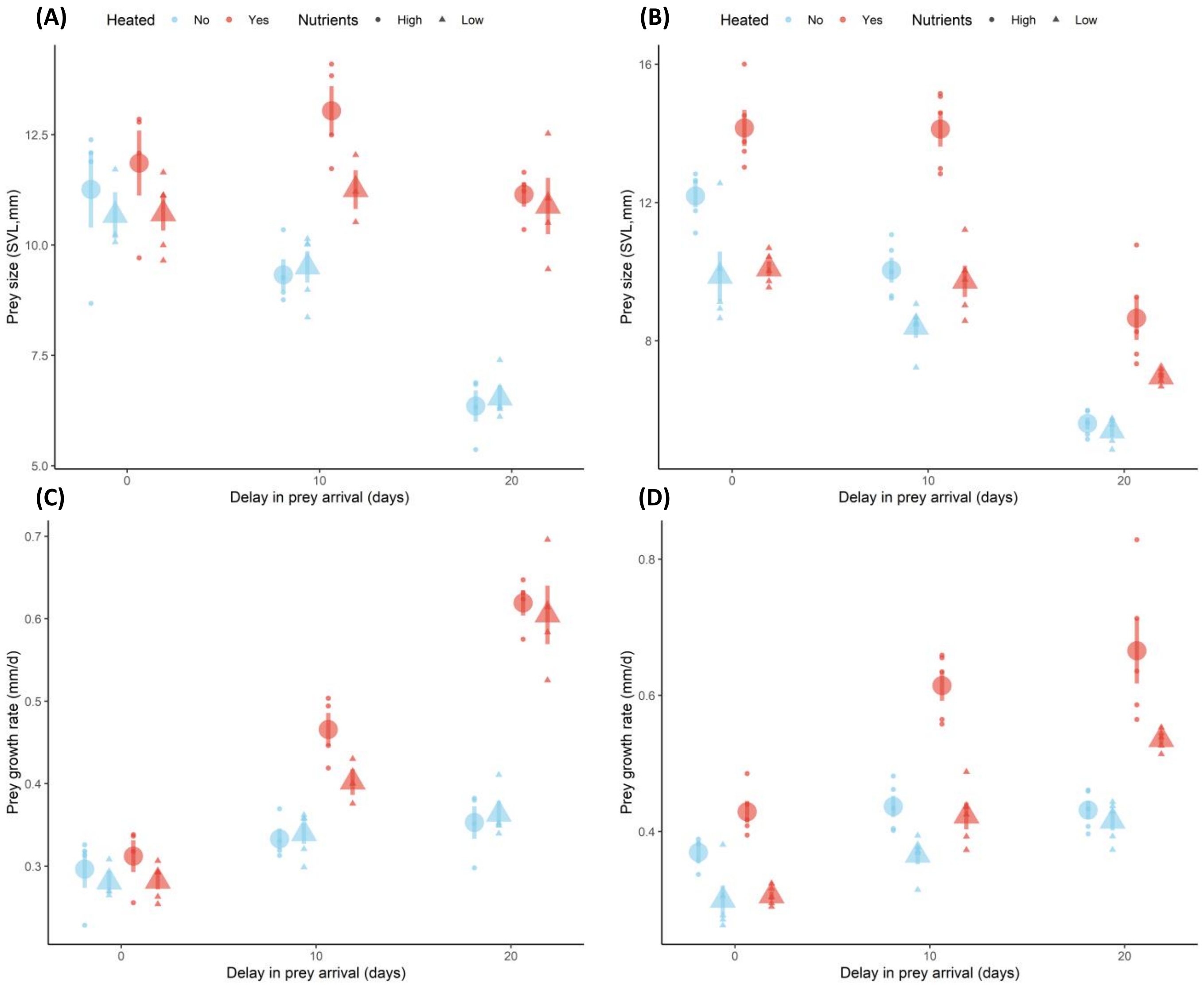

**Figure S3:** Mean mass, development time and number of emerging leopard frog metamorphs with dragonfly predators across arrival time, heating, and nutrient addition treatments. Large symbols indicate treatment mean ±1 SE, small symbols indicate mesocosm means. Symbols of different treatments are offset horizontally for a given arrival day treatment for visual clarity.

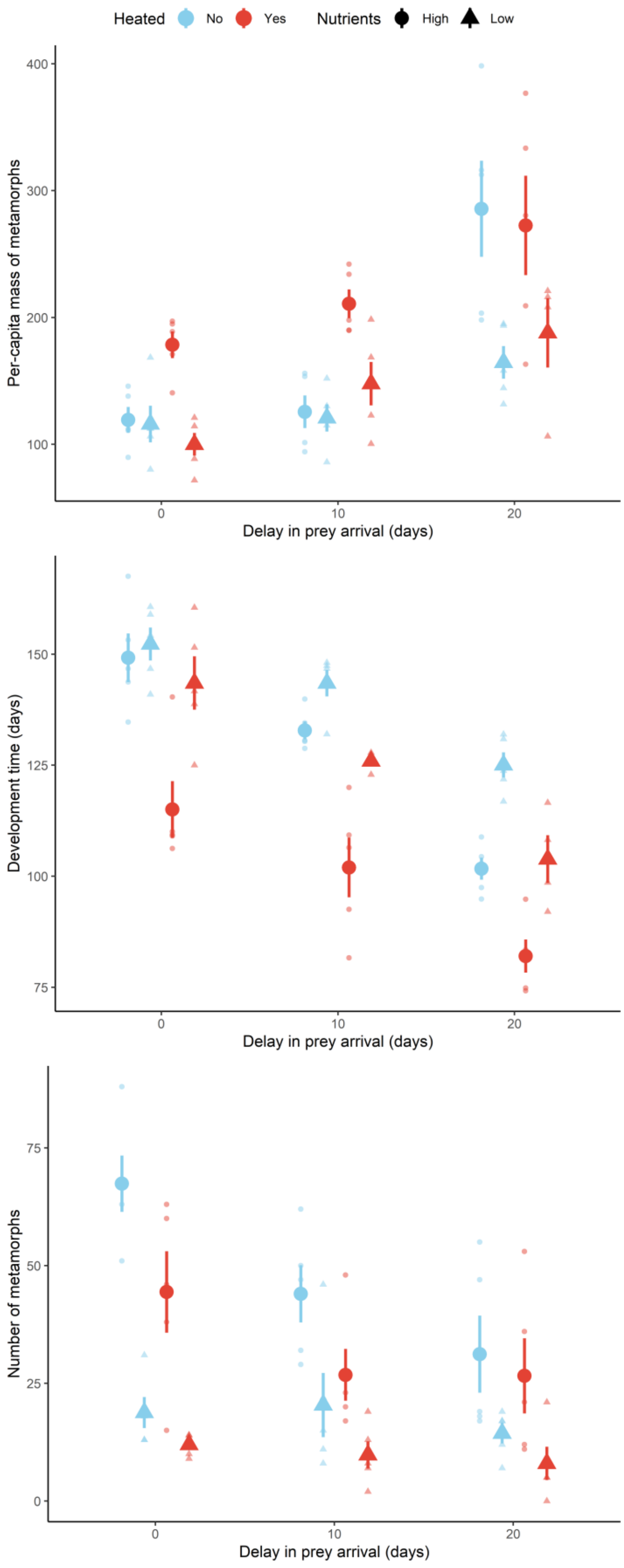

**Figure S4: Relationship between predator developmental time and prey survival across treatments.** Circles indicate given replicate using replicate means for predator development time, blue line indicates best fit linear model.

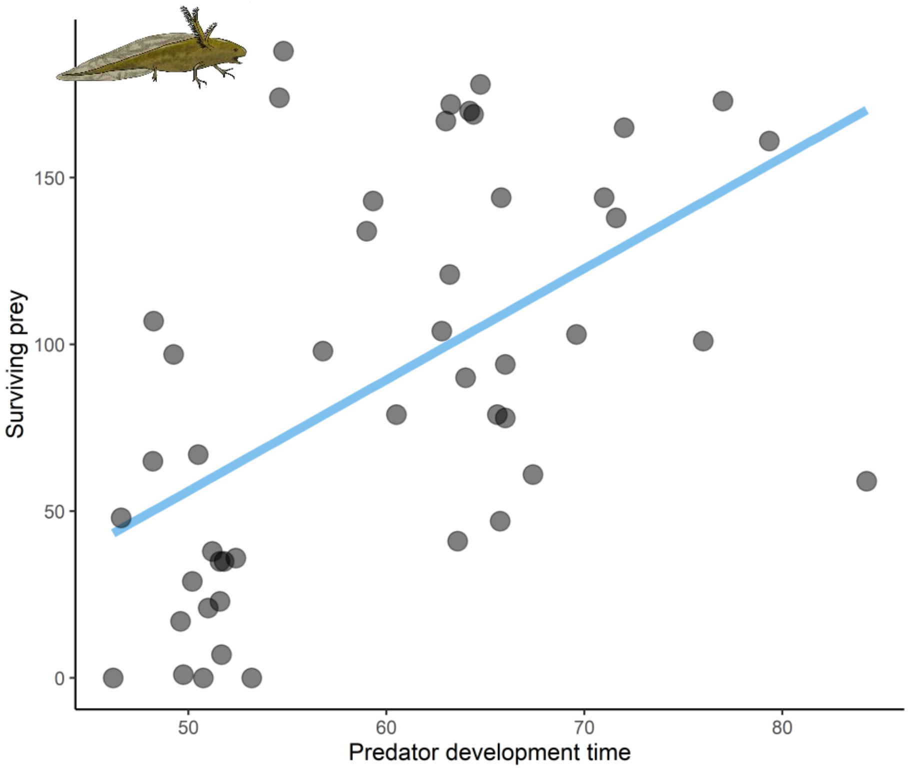


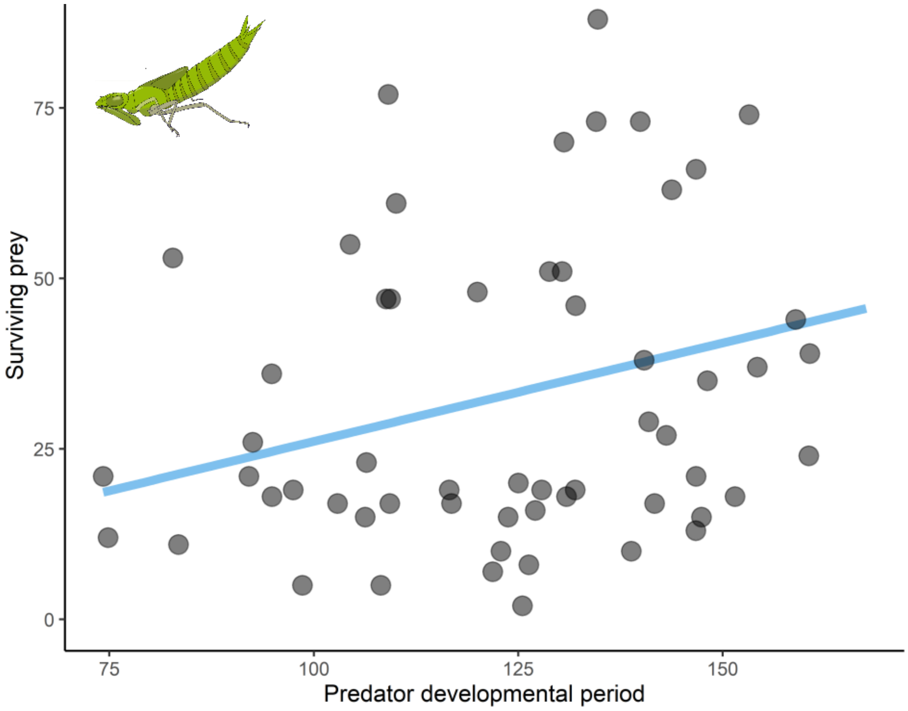

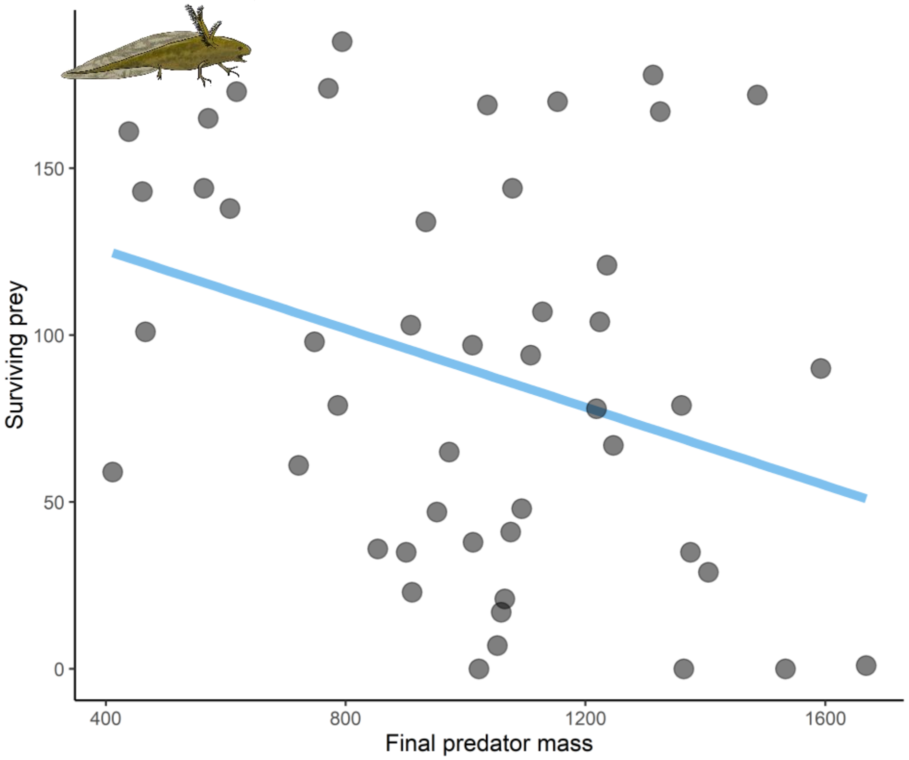

Surviving prey
150
100
50
0
400
800
1200
1600
Final predator mass

**Figure S5:** Change in periphyton (primary producer) over time for each prey arrival treatments (panels 0, 10, 20) depending on heating and nutrient additions. First row shows salamander predator system, second row dragonfly predator system.  Note difference in length of x axis between top and bottom panels.

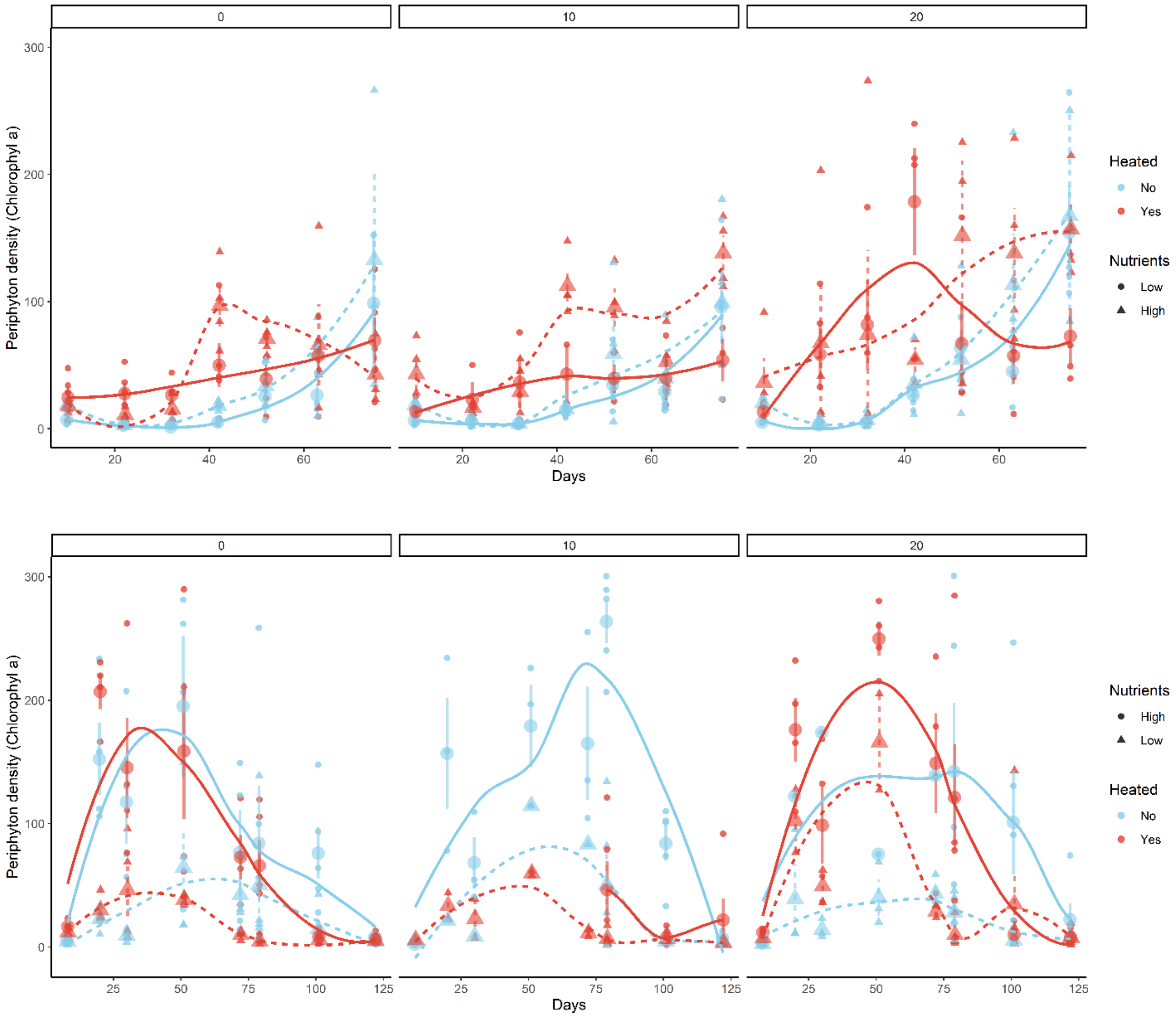
